## Supplementary figures and images for "The positionally conserved lncRNA DANCR is an essential regulator of zebrafish development and a human melanoma oncogene"

### Supplemental Figures

#### Supplemental Figure 1

A

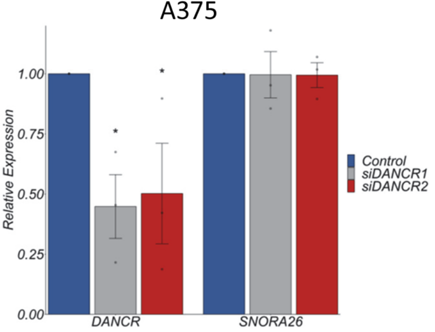

#### Supplemental Figure 2

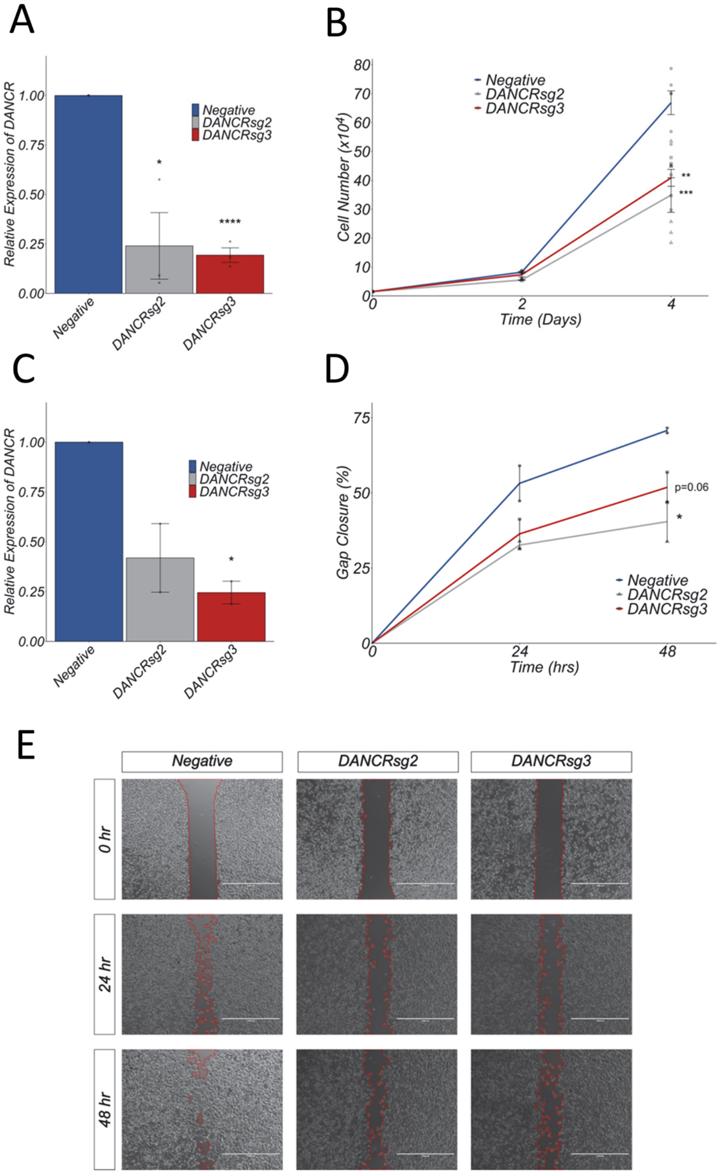

#### Supplemental Figure 3

A

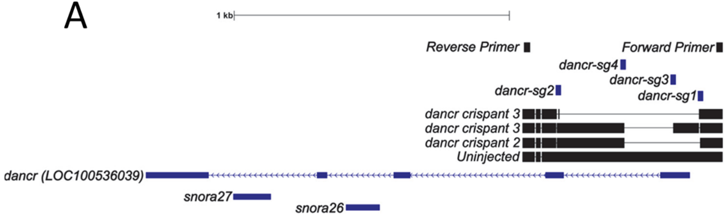


B

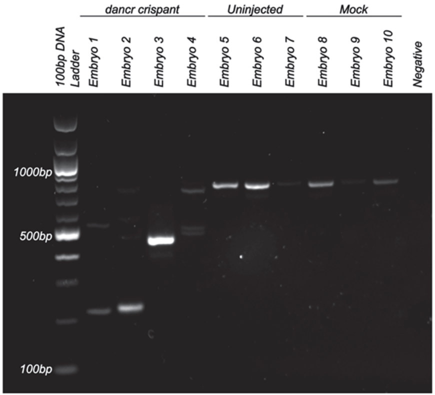
